# Sequence and phylogenetic analysis revealed structurally conserved domains and motifs in lincRNA-p21

**DOI:** 10.1101/2021.03.24.436769

**Authors:** Aditi Maulik, Devleena Bandopadhyay, Mahavir Singh

**Author notes:** To whom correspondence should be addressed, (A.M.); (M.S.).

## Abstract

Long Intergenic Non-coding RNAs (lincRNAs) are the largest class of long non-coding RNAs in the eukaryotes, which originate from the intergenic regions of the genome. A ~4kb long lincRNA-p21 is derived from a transcription unit next to the p21/Cdkn1a gene locus. LincRNA-p21 plays key regulatory roles in p53 dependent transcriptional repression and translational repression through its physical association with proteins such as hnRNP-K and HuR.It is also involved in the aberrant gene expression in different cancers. However, detailed information on its structure, recognition, and trans-regulation by proteins is not well known. In this study, we have carried out a complete gene analysis and annotation of lincRNA-p21. This analysis showed that lincRNA-p21 is highly conserved in primates, and its conservation drops significantly in lower organisms. Furthermore, our analysis has revealed two structurally conserved domains in the 5’ and 3’ terminal regions of lincRNA-p21. Phylogenetic analysis has revealed discrete evolutionary dynamics in these conserved domains for orthologous sequences of lincRNA-p21, which have evolved slowly across primates compared to other mammals. Using Infernal based covariance analysis, we have computed the secondary structures of these domains. The secondary structures were further validated by energy minimization criteria for individual orthologous sequences as well as the full-length human lincRNA-p21. In summary, this analysis has led to the identification of sequence and structural motifs in the conserved fragments, indicating the functional importance for these regions.

## Introduction

Long Intergenic Non-coding RNAs (lincRNAs) are one of the highly abundant and functionally important class of long non-coding RNAs (lncRNAs) in the eukaryotes, which originate from the intergenic regions in the genome (1). Although more than 3,000 human lincRNAs have been discovered using transcriptomic data and bioinformatics analysis (2–4), only a subset (less than 1%) have been functionally characterized, and determining the function of individual lincRNAs remains a challenge (5). Nuclear lincRNAs function by targeting chromatin-modifying complexes and regulating the transcription (5–7), whereas cytoplasmic lincRNAs regulate translational control of gene expression and transcript stability by binding with specific targets or by acting as competing endogenous RNAs (7–11). Few well-characterized examples are HOTAIR, Xist, and MALAT-1 lncRNAs that play roles in chromatin maintenance, X-chromosome inactivation, transcription regulation, cell motility, etc., showing that lincRNAs are a key regulator of diverse cellular processes (12–16). Several lincRNAs (e.g., MALAT1, H19) are also shown to be involved in carcinogenic processes by interacting with cancer-associated genes or other non-coding RNAs (17–21).

The functional diversities of lincRNAs arise from their ability to adopt different structures and molecular interactions with not only proteins but also with other RNAs and DNA (22, 23). LncRNAs are rapidly evolving, and the selection acting on structure rather than primary sequence has been proposed to explain their rapid evolution. This led to the “RNA modular code” hypothesis based on the view that evolutionary selection acts on structural domains in RNA (23–25). Some experimental evidence supports this concept. For example, the maternally expressed gene 3 (*MEG3*) lncRNA gene contains three distinct structure modules: M1, M2, and M3. Deletion analysis showed that motifs M2 and M3 are important for p53 activation. Intriguingly, a hybrid MEG3 transcript in which half of the primary sequence in the M2 motif was replaced by an entirely unrelated artificial sequence that displayed a similar secondary structure was fully functional in stimulating p53-mediated transcription (26). Similarly, for lincRNAs structural conservation rather than nucleotide sequence conservation seems to be crucial for maintaining their function (27). Several lincRNAs acquire complex secondary and tertiary structures, and their functions often impose only subtle sequence constraints.

Here, we have focused on the analysis of sequence, structure, and evolutionary features of lincRNA-p21 that is a transcriptional target of p53 and HIF1-a(28–31). LincRNA-p21 is a ~4kb long lncRNA derived from a transcription unit next to the p21/Cdkn1A gene locus (hence named lincRNA-p21) (6, 32). The ‘guardian of the genome’ tumor suppressor p53 plays a key role in maintaining genomic integrity (33). Upon DNA damage, p53 triggers a transcriptional response resulting in either cell cycle arrest or apoptosis (34). While p53 is known to transcriptionally activate numerous genes directly, the mechanism by which p53 causes gene repression involves its interaction with other factors. For example, several lincRNAs that are physically linked with repressive chromatin-modifying complexes have been shown to act as repressors in p53 transcriptional regulatory networks (5). LincRNA-p21 has been identified as one among such p53-activated lincRNAs (29). p53 binds in the highly conserved canonical p53-binding motifs containing promoter regions of lincRNA-p21, thereby driving its expression (35, 36). LincRNA-p21, in turn, functions as a downstream transcriptional repressor of several genes hence plays an important role in the p53-dependent induction of cell death in response to DNA damage. In recent times, numbers of reports have shown misregulation and differential expression of lincRNA-p21 in a number of cancers, including prostate, colorectal, chronic lymphatic leukemia, and atherosclerosis (37–41).

LincRNA-p21 mediates the transcriptional repression through specific association with heterogeneous nuclear ribonucleoprotein K (hnRNP-K). hnRNP-K was previously identified as a component of the repressor complex that acts in the p53 pathway (42, 43). It has the ability to bind both ssDNA and ssRNA via its three KH (hnRNP-K homology) domains (44). Furthermore, lincRNA-p21 was shown to bind hnRNP-K through a conserved 780 nucleotides long 5’ region. Interaction of hnRNP-K with lincRNA-p21 was shown to be required for its proper localization and subsequent induction of apoptosis via transcriptional repression of p53-regulated genes (29). Overall, these studies implicated lincRNA-p21 – hnRNP-K association to mediate the p53 dependent transcriptional repression of several genes.

LincRNA-p21 has been shown to be transported to the cytoplasm, where it is also involved in repressing the translation of several mRNAs (for example, JUNB, β-catenin, andpotentially many more that encode proteins involvedin cell proliferation and survival). LincRNA-p21 is hypothesized to directly base-pair with mRNAs and in conjunction with knowntranslational repressor RCK/p54 (the evolutionarilyconserved ATP-dependent DEAD-BOX helicase)remodel mRNPs and influence their association with elongation-factor eIF4E. The levels oflincRNA-p21 in the cytoplasm are regulated by RRMdomains containing the proteinHuR/ELAVL1 (7). The association of HuR with lincRNA-p21 favored the recruitment of let-7/Ago2 to lincRNA-p21, leading to the lower lincRNA-p21 stability. Thisleads to relieving of translation repression of lincRNA-p21 targeted mRNAs (7, 45).

In general, lncRNAs show poor sequence conservation across species showing only scattered conserved regions surrounded by large seemingly unconstrained sequences (27). Therefore, phylogenetic analysis of lncRNAs is important and can reveal the conserved and functionally important regions in RNA. A functional domain in lncRNA is likely to be conserved in animals and adopt a structure that is possibly shared by the orthologues sequences. This pattern of conservation allows computational genome analysis that can be used to identify lncRNAs from different organisms as well as define their functional domains. This approach had been adopted to understand the origin and evolution of lincRNAs such as XIST and HOTAIR (46–48). In this study, using computational and bioinformatic methods, we have investigated lincRNA-p21 that function in both cis and trans to understand its sequence, structure, and evolution (28, 29). Particularly, we addressed the following questions. First, whether lincRNA-p21 exists and shows evolutionary conservation in all mammals or vertebrates. To address this question, we looked into 13 vertebrate genomes using Infernal, (INFERence of RNA ALignmen) a structure-based RNA homology search program (49). Our results showed that orthologous sequences of lincRNA-p21 exist only in mammals. Infernal hits were found well conserved in closely related species but poorly conserved among all other animals. Second, evolutionarily conserved features of lincRNA-p21 were investigated using PAML and EvoNC computational programs by analyzing the sequences orthologous to lincRNA-p21 (50). LincRNA-p21 conserved domains showed discrete evolutionary dynamics, with different nucleotide substitution rate amongst different mammals. Additionally, lincRNA-p21 has undergone accelerated evolution compared to the neighboring protein-coding Cdkn1a gene. Finally, using PMmulti (51) and Mfold (52) algorithms, we have predicted the secondary structures of conserved regions of lincRNA-p21 from different animals and also a full lincRNA-p21 sequence. This analysis revealed invariable fragments in these structures, which may have functional roles in lincRNA-p21’s functions.

## Methods

### Data

The sequence of human lincRNA-p21 long isoform (accession number KU881768.1) and mouse lincRNA-p21 (accession number NC_000083.6) were acquired from the National Center for Biotechnology Information (NCBI) database. The unmasked genome data of a human (GRCh38, Dec. 2013), chimpanzee (Pan_tro_3.0, May. 2016), rhesus monkey (Mmul_8.0.1, Nov. 2015), gorilla (gorGor4, Dec. 2014), cow (ARS-UCD1.2, Apt. 2018), horse (EquCab3.0, Jan. 2018), dolphin (turTru1, Jul. 2008), cat (Felis_catus_9.0, Nov. 2017), mouse (GRCm38, Jan. 2012), rat (Rnor_6.0, Jul. 2014), platypus (OANA5, Dec 2005), chicken (GRCg6a, Mar 2018), and zebrafish (GRCz11, May. 2017) were downloaded from Ensembl (release 97).

### Acquiring sequences orthologous to mouse lincRNA-p21 exons through genome search

Human and mouse lincRNA-p21 sequences were aligned in Emboss (53). The human lincRNA-p21 sequence ranging from 1 to 198 nucleotides and 199 to 3898 nucleotides were found to align with mouse lincRNA-p21 exon 1 and exon 2 sequences, respectively (Figure 1). Accordingly, for the purpose of our study, the human lincRNA-p21 sequence was divided into two segments: 1 – 198 and 199 – 3898 nucleotides sequence spans. These two consecutive sequence ranges in human lincRNA-p21 were considered as mouse equivalent of human exon 1 and exon 2, respectively, and termed as segment 1 and segment 2 for human lincRNA-p21 (Figure 1). Each of these two segments was separately used as a query to search the genomes of the chimpanzee, rhesus monkey, gorilla, cow, horse, dolphin, cat, mouse, rat, platypus, chicken, and zebrafish in Ensembl using BLASTN (54). For each query, the best hit obtained in a particular genome was considered as the sequence orthologous of human lincRNA-p21 for the corresponding species. Likewise, the obtained sequence orthologous in humans, rhesus monkey, and cat were aligned using LocARNA (55). Based on these alignment results, two queries (query 1 and query 2) were built using cmbuild and cmcalibrate functions of Infernal [v1.1] (49). Cmbuild is a program that builds a covariance model from an input multiple alignment and cmcalibrate calibrates E-value parameters for the covariance model. The calibrated models were used to search the whole genomes of 13 vertebrates using the cmsearch function of Infernal, a program that searches for a covariance model against any sequence database.

**Figure 1.**
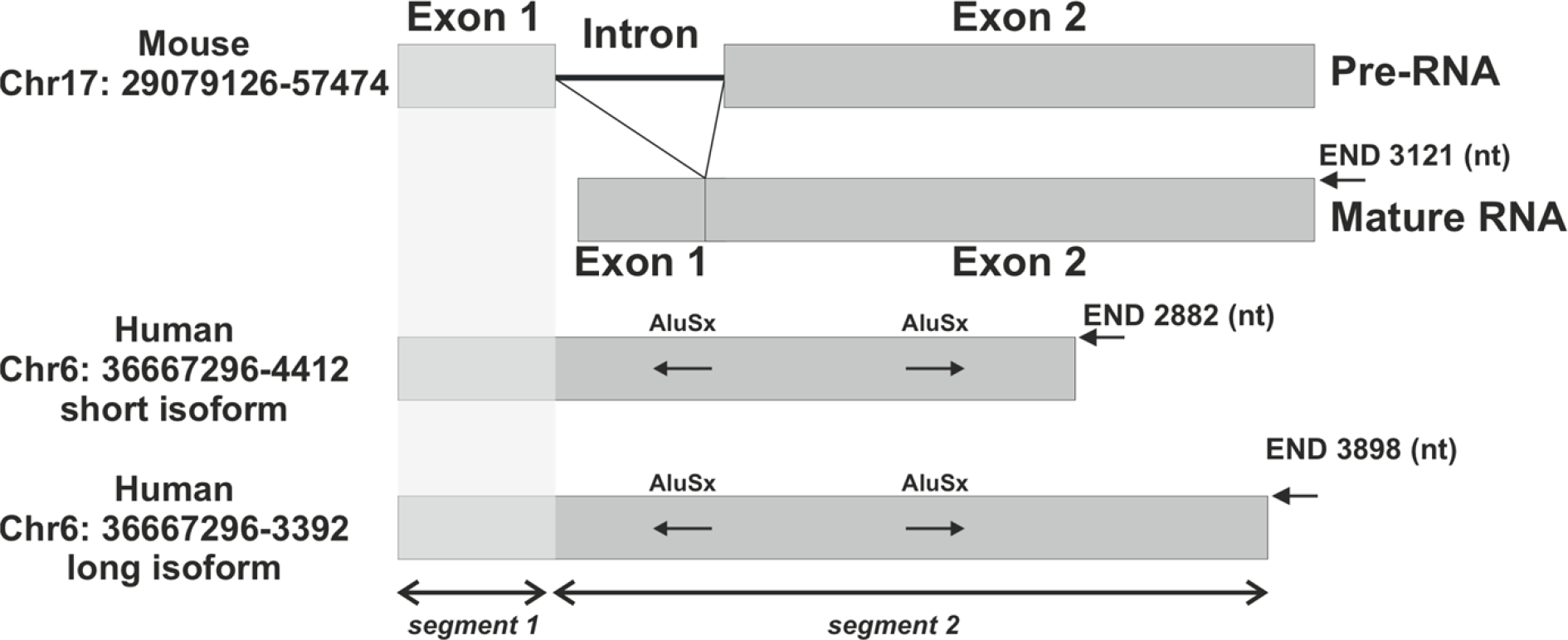
Schematic of precursor (Pre-RNA) and mature mouse lincRNA-p21 (2 exons lncRNA), short and long isoform of human lincRNA-p21 (single exon lncRNA). The shaded box represents a region of sequence homology between mouse and human lincRNA-p21 (both SIsoE1 and LIsoE2). The Alu repeats found in the human lincRNA-p21 isoforms are mentioned. The two segment spans, 1 – 198 and 199 – 3898 nucleotides in human lincRNA-p21 sequence were also marked.

### Structure prediction through sequence alignment and using a thermodynamic method

Sequences that correspond to the highest-scoring Infernal hits of the ten mammals were considered as orthologous to the two lincRNA-p21 segments and named as domain A and domain B. They were aligned using cmalign function of Infernal for phylogenetic analysis. Structures were predicted for the orthologous of these two domains using PMmulti (51) and Mfold (52). PMmulti performs pairwise and multiple progressive alignments of RNA sequences, and Mfold predicts the secondary structure of RNA and DNA, mainly by using thermodynamic methods. Predicted structures were displayed using either Mfold (52) or PseudoViewer (56). In all cases, default parameters were used.

### Phylogenetic study

Using orthologous sequences of lincRNA-p21 domain A and domain B in the 10 mammals, two phylogenetic trees were built with 500 bootstraps using the DNAdist and Kitsch functions of Phylip (the Phylogeny Inference Package) (Felsenstein J: Phylip: Phylogenetic inference program. Version 3.6 University of Washington Seattle; 2005). DNAdist calculates four different distances between species using nucleic acid sequences. The distances can then be used in the distance matric based phylogeny estimation program Kitsch. We performed a phylogenetic analysis of the two trees using the baseml function of PAML (Phylogenetic Analysis by Maximum Likelihood v4.4) (57). Baseml is for Maximum Likelihood (ML) phylogeny analysis of nucleotide sequences that estimates tree topology, branch lengths, and substitution parameters under a variety of nucleotide substitution models. Fixed parameters included model = 4 (the HKY85, Hasegawa-Kishino-Yano, 85 nucleotide substitution model); fix_kappa = 0 and kappa = 2; fix_alpha = 0 and alpha = 0.5; ncatG = 5, fix_rho = 1 and rho = 0; and cleandata = 0. The parameters kappa (κ, the transition/transversion rate ratio), alpha (α, shape parameter of the gamma distribution), local clock, and rates of substitution (r_1_, r_2_ and r_3_) were estimated under different conditions (58–61). The evolution of the conserved domain A and the neighboring Cdkn1a gene was analyzed in 10 mammals using EvoNC (50). For this purpose, the rate of substitution in noncoding regions relative to the rate of synonymous substitution in coding regions is estimated by a parameter ζ. ζ is the nucleotide substitution rate in the noncoding region, normalized by the synonymous nucleotide substitution rate in the coding region. Therefore, when a site is subject to neutral selection ζ = 1. Similarly, ζ > 1 indicates positive selection, while ζ < 1 suggests the presence of negative selection. Therefore, the interpretation of ζ is similar to the interpretation of the rate of nonsynonymous/synonymous substitution (ω) in models of evolution in coding regions.

## Results and Discussion

### The sequences of lincRNA-p21 orthologs show poor conservation among vertebrates

LincRNA-p21 is located between Srsf3 and Cdkn1a protein-coding genes on chromosome 6 in human. In mouse, the mature lincRNA-p21 is found on chromosome 17, consists of two exons: exon 1 and exon 2 while, human lincRNA-p21 contains only a single exon on chromosome 6 that aligns with both exon 1 and exon 2 of mouse lincRNA-p21 (Figure 1). We searched several vertebrate genomes in the UCSC genome browser for matches to lincRNA-p21. The whole sequence of human lincRNA-p21 showed apparent conservation among mammalian orthologs (Figure 2A and 2B). Individually, when mouse exon 1 and exon 2 equivalent human lincRNA-p21 sequences, (defined as segment 1 and segment 2 respectively) were searched using BLAT and BLASTN (62) tool across different vertebrate genomes in Ensembl databases, they returned several hits with high to low scores. We assumed that if a hit is found in between Srsf3 and Cdkn1a genes with an E-value cut off 1e-05, it can be considered as orthogous to human lincRNA-p21 sequence. Notably, for both the queries, the hits produced with highest score (least E-value) were always located between the two proteincoding genes for all the different mammalian genome studied here. This result suggests that lincRNA-p21 has orthologs in mammals. The top scoring hit from each genome are listed in Table 1. The hits found in primates were high scoring but hits from all other non-primate mammals had comparatively poor scores. In other words, close matches were observed only in primates and not in other mammals. Eventually, for non-mammalian vertebrates, like platypus, chicken and zebrafish no hits were found between the two protein coding genes. This finding implies that if lincRNA-p21 also has orthologs in non-mammalian vertebrates, they may have moderate to low sequence conservation to be identified or revealed solely by sequence search method. The compensatory mutations commonly found in ncRNAs that changes the primary sequence but maintains the secondary structures can be one of the reasons behind poor sequence conservation (63, 64).

**Figure 2.**
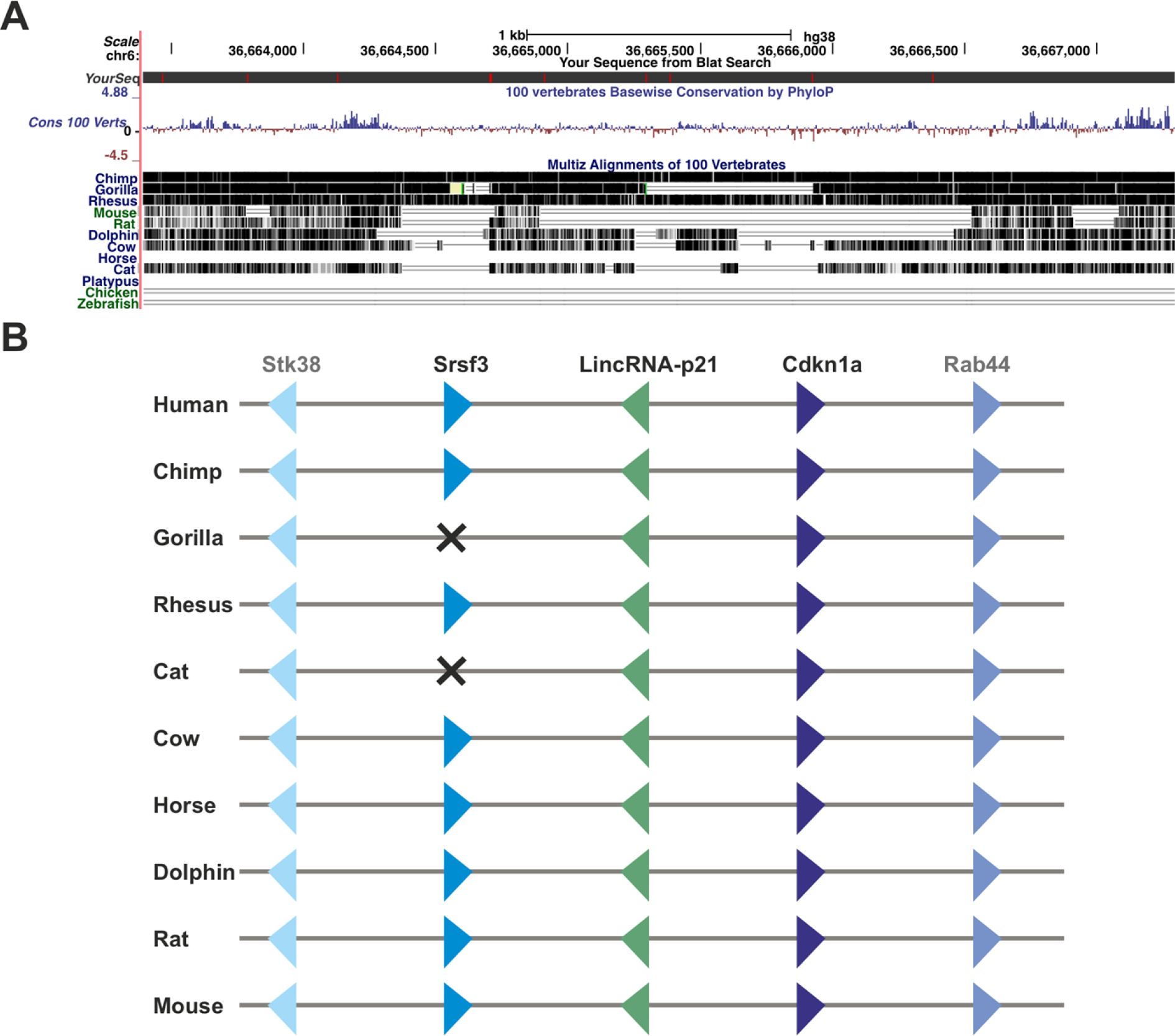
Sequence conservation of lincRNA-p21 orthologues in mammals. (A) The sequences of lincRNA-p21 orthologous are conserved in primates but less conserved in other animals (from UCSC Genome browser). (B) The order and orientation of lincRNA-p21 and its neighboring protein coding genes in mammals (X represents the absence of Srsf3 gene).

**Table 1.** BLASTN result from Ensemble genome browser

| Ensemble Hits |  |  | Segment 1 |  | Segment 2 |  |
| --- | --- | --- | --- | --- | --- | --- |
|  |  |  | Score | E-value | Score | E-value |
| Mammals | Primates | Human (chr6) | 392 | 5e-106 | 7198 | 0.0 |
|  |  | Chimp (chr6) | 392 | 3e-107 | 6784 | 0.0 |
|  |  | Gorilla (chr6) | 392 | 3e-107 | 6597 | 0.0 |
|  |  | Rhesus monkey (chr4) | 336 | 3e-90 | 5174 | 0.0 |
|  | Ungulates | Cow (chr23) | 73.8 | 3e-11 | 246 | 7e-62 |
|  |  | Horse (chr20) | 71.9 | 1e-10 | 174 | 2e-40 |
|  |  | Dolphin | 67.9 | 2e-09 | 281 | 1e-72 |
|  | Carnivorous | Cat (chrB2) | 60 | 4e-07 | 174 | 3e-40 |
|  | Rodents | Rat (chr20) | 61.9 | 1e-07 | 176 | 6e-41 |
|  |  | Mouse (chr17) | 65.7 | 8e-09 | 152 | 1e-33 |
|  | Ancestral mammal | Platypus | No hit found with Srsf3 and/Cdkn1a |  |  |  |
| Other vertebrates | Bird | Chicken | No hit found with Srsf3 and/ Cdkn1a |  |  |  |
|  | Fish | Zebrafish | No hit found |  | No hit found with Srsf3 and/ Cdkn1a |  |

### LincRNA-p21 exists in mammal and show high conservation among primates

LncRNAs are characterized not only by varying sequences but also conserved structures. Therefore, we extended our investigation through structural search in different vertebrate genomes to confirm the existence of lincRNA-p21 only in mammals. Infernal was used to search the whole genomes for matches to lincRNA-p21. Infernal is a local RNA alignment and search program that uses the combination of sequence consensus and secondary structure conservation in RNA to generate a covariance model structure (49). To construct the query for building covariance model of lincRNA-p21 (necessarily a representative structure), we used human lincRNA-p21 segment 1 and segment 2 corresponding sequences and their identified orthologous sequences in rhesus monkey (BLASTN score: 336, E-value: 3e-90 and BLASTN score: 5174, E-value: 0, for segment 1 and 2, respectively), a primate that is distantly related to human and cat (BLASTN score: 60, E-value: 4e-07 and BLASTN score: 174, E-value: 3e-40 for segment 1 and 2, respectively) which is a non-primate mammal. Two queries (query 1 and query 2) were built with these sequences using the cmbuild and cmcalibrate functions of Infernal tool (49). These two queries were used to search the complete genome of ten placental mammals (human, chimpanzee, rhesus monkey, gorilla, cow, horse, cat, dolphin, mouse, and rat), the ancestral mammal platypus, and two other vertebrates (non-mammals) chicken and zebrafish (Figure 3). Orthologous sequences of the lincRNA-p21 segments (hits located between Srsf3 and Cdkn1a with high scores) were obtained through Infernal search in all of the placental mammals but not in platypus or the other non-mammalian vertebrates (Figure 3). Notably, each query produced just one high scoring hit in the mammalian genomes (Table 2). All other hits had low Infernal (cmsearch) scores. For example, in gorilla genome, the score from the top hit to the next hit had drastically reduced from 246.2 (E-value 3.9e-66) to 23.7 (Evalue 0.53) for query 1 and from 202.4 (E-value 6.5e-43) to 57.9 (E-value 9.4e-08) for query 2. Infernal score for the top hits for query 1 ranged from 246.2 (E-value 3.9e-66) in gorilla to 84.8 (E-value 2.9e-18) in mouse genome. Similarly, for the query 2, values for the top hits ranged between 203.6 (3.5e-43) in rhesus monkey to 71.4 (3.9e-11) in horse. These hits were located between Stk38/Srsf3 and Cdkn1a/Rab44 (note: Srsf3 is absent in some mammals, Figure 2B). Overall, query 1 did not produce any high-scoring hits in rodents. Similarly, query 2 did not result in any high-scoring hits in ungulates like horse or dolphin. Moreover, both the queries produced good matches in primates but poor matches in the remaining mammals (Figure 3 and Table 2).

**Figure 3.**
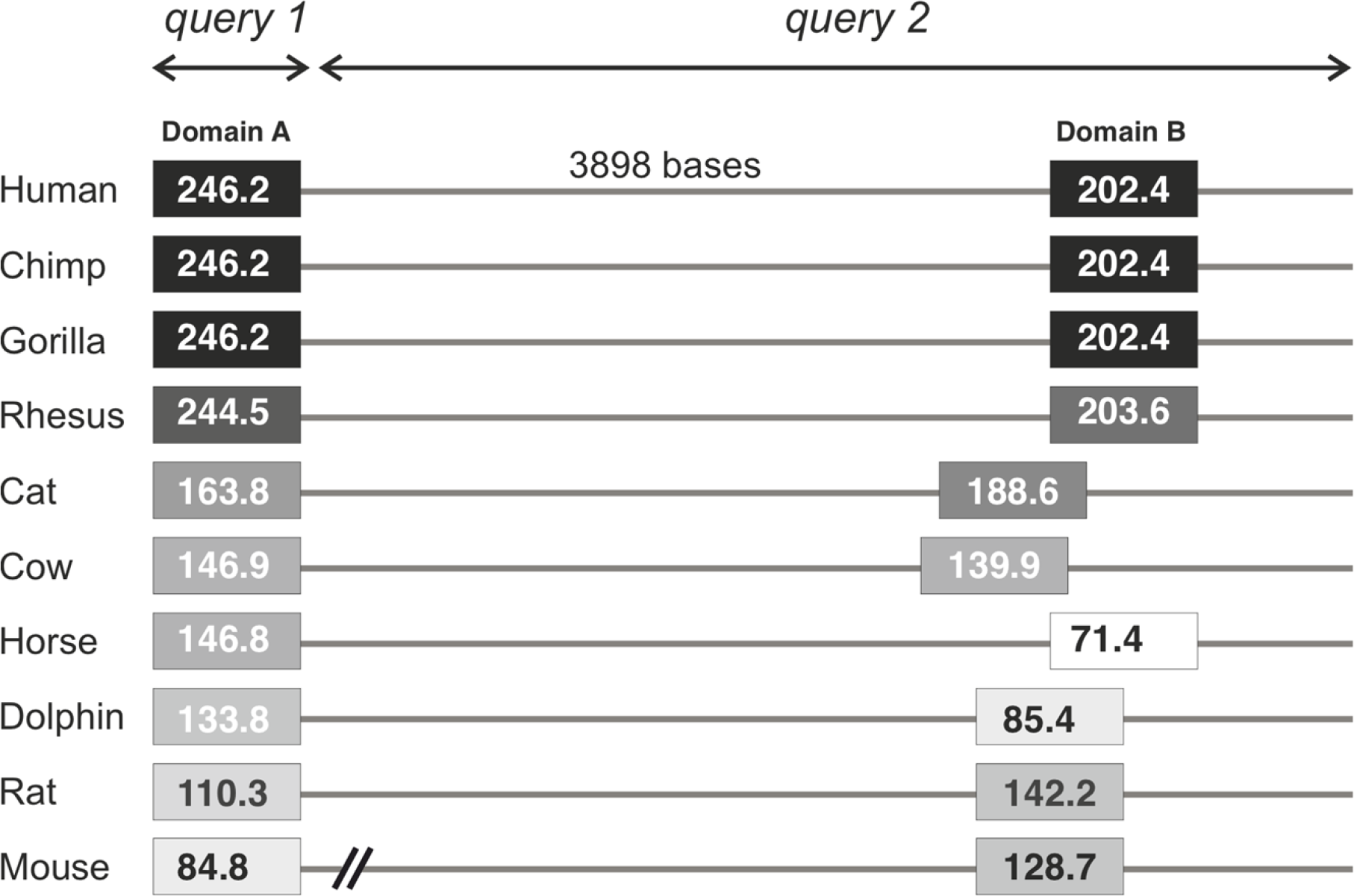
Orthologous of lincRNA-p21 exists only in mammals. Depiction of Infernal search result where each grey box represents hit with highest score (scores are written in the boxes) found for the respective genomes. Better conservation (better scores) is indicated by the darkness of the boxes. In each of the searched genomes two conserved domains are found for linRNA-p21, domain A (left side): at 5’ terminal region that spans for the entire Query 1 and domain B (right side): at 3’-terminal region, much shorter fragment than the Query 2. Black lines indicate the unmatched part in the Infernal search. The double slashes in the schematic of the mouse gene indicate long introns.

**Table 2.**
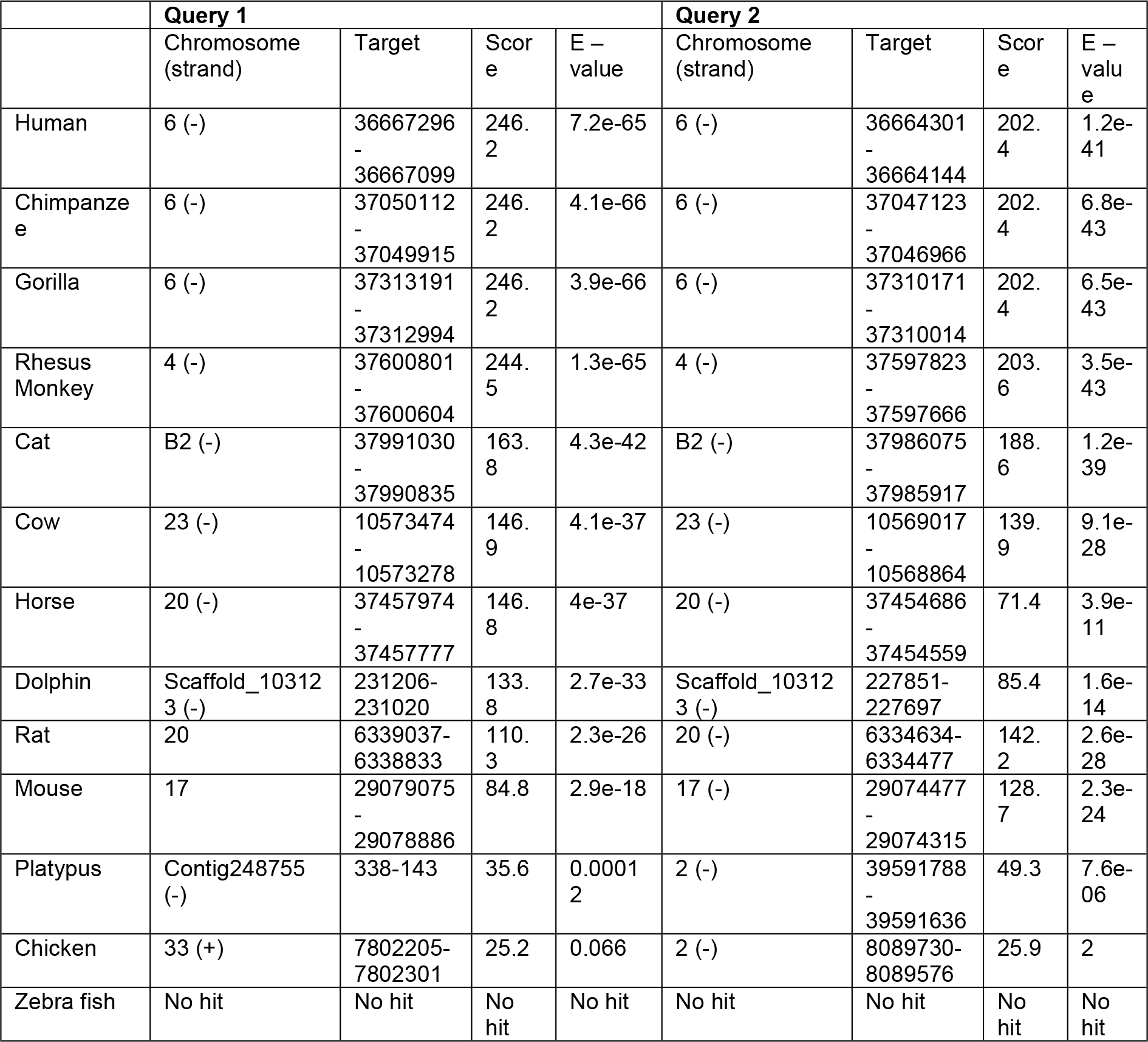
Description of Infernal hits with high scores and successive addresses in individual genome using Query 1 and Query 2

However, the top hits from non-mammalian vertebrates (platypus, chicken, zebra fish) had much less scores compared to the mammalian species (Table 2). For example, in chicken, the highest scoring hit had Infernal score as less as 25.2 (E-value 0.0666) and 25.9 (E-value 2.0) for query 1 and query 2, respectively. Moreover, the hits were not found between Srsf3 and Cdkn1a genes. Thus, combination of sequence and structural search confirmed that lincRNA-p21 exists only in mammals and it has evolved further to become highly conserved in primates.

We hypothesize that lincRNA-p21 may have conserved structural domains but divergent sequences in different vertebrates. This is a general feature for lncRNAs (65). For example, Xist and HOTAIR both contain fast evolving sequences as well as highly conserved structures (48, 66). The reasons that constrain lncRNAs evolution are not clear. Understanding of evolutionary constraint of lncRNA like lincRNA-p21 that functions both in cis and trans to control local and global gene expressions will be more intriguing.

### Two conserved domains identified in the 5’ and 3’ terminal regions were found unique to lincRNA-p21

Besides one high-scoring hit located between Srsf3 and Cdkn1a, several scattered low-scoring hits of queries (query 1 and 2) were widely obtained in mammalian as well as other vertebrate genomes. In different genomes, number of non-specific hits varied across human, chimpanzee, gorilla, rhesus monkey, cow, horse, dolphin, rat, mouse, platypus, chicken and zebra fish for both query 1 and query 2. The 3707 nucleotides long query 2 produced more numbers of low-scoring hits compared to 198 nucleotides long query 1 that received far less numbers of hits. However, whether these hits have any functional roles is not clear. These low-scoring hits are expected to be insignificant and random. Since along with lincRNA-p21, a few other lncRNAs were also reported to interact with polycomb proteins, Infernal search might have detected some of those consensus sequences for the protein binding shared by different lncRNAs. Considering that the best hit was less conserved in non-primate mammals and much shorter in all other genomes (Figure 3), we inferred that for query 2 functionally conserved domain(s) in mammals must be much shorter than 3707 nucleotides. However, as the best hits for query 1 spanned the entire length for all the mammals (Figure 3), we assume that entire region of 198 nucleotides long segment 1 may have a structurally conserved function. The data in Figure 3 show that the best structural hits were found as shorter fragments one at the 5’-terminal region (domain A) and another towards the 3’-terminal region (domain B) of full-length lincRNA-p21. Result of single high scoring hit, found in each case also suggests that these two domains in segment 1 and segment 2 (henceforth called domain A and domain B respectively) could form the unique, minimum functional structural unit of lincRNA-p21, which is not shared by other lncRNAs. Thus, we conclude that the functional domain(s) of lincRNA-p21 conserved in mammals should be much shorter than ~4 Kb length of this RNA. The orthologous of domains A and B were conserved in primates with Infernal score (E-value) ranging from 246.2 (3.9e-66) to 244.5 (1.3e-65) and from 203.6 (6.5e-43) to 202.4 (3.5e-43), respectively and were less conserved in other mammals with 163.8 (4.3e-42) to 84.8 (2.9e-18) and 188.6 (1.2e-39) to 71.4 (3.9e-11) Infernal score. Furthermore, while the orthologous of domain A were found to be much less conserved in rodents with Infernal score of 110.3 (2.3e-26) and 84.8 (2.9e-18) for rat and mouse (Figure 3), the orthologous of domains B were less conserved in ungulates like dolphin and horse with 85.4 (1.6e-14) and 71.4 (3.9e-11) Infernal score, respectively. This observation suggests that the two domains have experienced different evolutionary constrains.

### Phylogenetic distribution of orthologous sequences of the conserved domains of lincRNA-p21

Protein coding genes commonly originate by gene duplication followed by neofunctionalisation (67–69) and/or subfunctionalisation (70, 71). However, the process and dynamics of evolution in non-coding RNAs is not well understood. We decided to analyze the molecular evolution of the lincRNA-p21 in detail. Using the sequences orthologous to domain A and the sequences orthologous to domain B, two phylogenetic trees were built using Phylip (Figure 4 and also please see Methods section). We assumed that nucleotide substitutions followed the HKY85 nucleotide substitution model (58) and rates of nucleotide substitutions varied among sites, to investigate orthologous sequences of lincRNA-p21 using PAML (http://abacus.gene.ucl.ac.uk/software/paml.html).We compared both the trees under a condition in which nucleotide substitution rates were variable among sites and the molecular clock was allowed to vary from branch to branch. In such a situation, tree in Figure 4A estimated a slightly smaller log-likelihood (−805.4 vs. −808.15) for orthologous of domain A but slightly larger log-likelihoods for the other orthologous set (−697.02 vs. −700.25) in Figure 4B. We checked whether nucleotide substitution rates varied among sites in each of the domains using the log-likelihood ratio test (a statistical test for comparing two models) (61). Similar value of 2ΔlnL = 2((−805.4)-(−808.15)) = 5.5 and 2((−697.02)-(−700.25)) = 6.46 were obtained from orthologous sequences of domain A and domain B, respectively (HKY85+gamma model called the HKY85 model). The probability distribution of the test can be approximated by a Chi-square distribution with one degree of freedom, with χ^2^_1_, _2_% = 5.41, supporting the model of variable nucleotide substitution rates. Our analysis also revealed that the orthologous sequences of the two domains had different transition/transversion rate ratio (κ) (Table 3). Because both the domains had α (shape parameter of the gamma distribution for variable substitution rates across sites (72)) value >1 (Table 3), most sites in these regions should undergo moderately high substitution rates, although a few sites might have slow rates of substitution.

**Figure 4.**
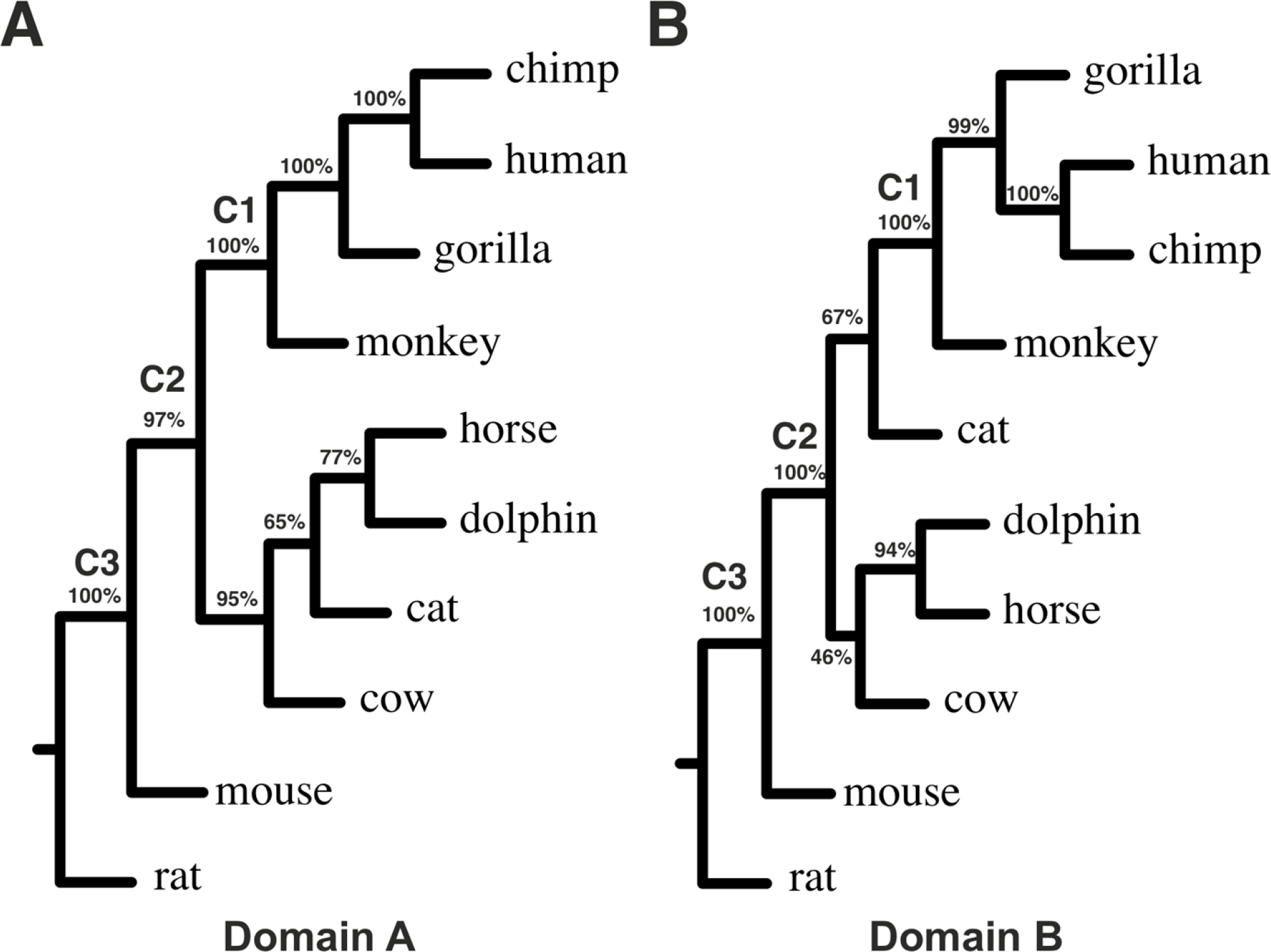
Phylogeny of lincRNA-p21. (A) Tree built with sequences of orthologous of domain A and (B) domain B. C1, C2 and C3 indicate three local clocks inserted at three different places for different computations. Bootstrap values are also mentioned at each node.

**Table 3.** Estimated parameters of molecular evolution from PAML.

|  | Domain A | Domain B |
| --- | --- | --- |
| Kappa ($\kappa$) | 3.56 | 2.65 |
| Alpha ($\alpha$) | 1.35 | 1.34 |
| $r_1/r_2$ | 2.26<br>4.88 | 2.15<br>8.0 |
| $r_1/r_3$ | 5.7<br>8.82 | 0.81<br>1.81 |
| Kappa ($\kappa$) and Alpha ($\alpha$) refer transition/transversion rate ratio and shape parameter of the gamma distribution, respectively. $r_1$, $r_2$ and $r_3$ denote different nucleotide substitution rates between clades. | | |

To investigate lincRNA-p21 evolution in further detail, we explored whether nucleotide substitution rates varied among clades. A log-likelihood ratio test was performed to determine whether the HKY85 model would fit the data better with or without a global clock. 2ΔlnL = 2((−805.4)-(−821.75)) = = 32.7, was obtained from orthologous sequences of domain A (−805.4 for the HKY85+gamma model without a global clock and −821.75 for the HKY85+gamma model with a global clock). Similarly, this value is 81.62 (2((−697.02)-(−737.83)) = 81.62) for domain B. This log-likelihood ratio test has eight degrees of freedom. Therefore, the probability distribution of the test approximated by the Chi-square distribution with eight degrees of freedom, χ^2^_8, 2_% = 18.17, negated the global clock hypothesis for both the domains. Then, we set three local clocks (C_1_, C_2_ and C_3_) inserted at three different places in the trees to determine whether the domains evolved at different rates among mammals. To obtain stable local clock estimations, only two local clocks (C_1_, C_2_ and C_1_, C_3_) were specified in each computation and the remaining species had rate equals to 1 (r_0_). For both the domains, the substitution rates (Table 3) estimated for primates (r_1_) were low compared to non-primate mammals (r_2_ and r_3_). Table 3 shows the rate of evolution, r_2_ in ungulates and carnivores (4.88) is more than twice of r_1_ in primates (2.26) for domain A. Similarly, for domain B, r_2_ in ungulates and carnivores (8.0) is more than 3 times than r_1_ in primates (2.15). Similar trends were observed for both Domain A and domain B while computing rate of evolution, r_1_ in primates vs. r_3_ that includes rodent for local clock estimation. Therefore, these varied rates of nucleotide substitution among clades suggested lincRNA-p21 domains followed discrete evolutionary dynamics in mammals with slow evolution rate in primates. Interestingly, a 5’ segment of lincRNA-p21 has been reported to bind to the hnRNP-K protein (29), thus whether the slow rate of evolution of domain A in primates indicate any relationship with its protein binding functions needs further investigations.

### LincRNA-p21 is rapidly evolving compared to its nearby protein coding Cdkn1a gene

Cdkn1A and Srsf3 are the two neighboring protein coding genes of lincRNA-p21. Because Srsf3 is absent in gorilla and cat, we compared the evolution of the exon of Cdkn1a gene along with the evolution of the domain A of lincRNA-p21 in ten mammals (see Methods section for details). The Cdkn1a genes exist commonly in all vertebrates, unlike the lincRNA-p21 gene exists only in mammals. Therefore, we were interested to know whether lincRNA-p21 has evolved faster or slower than the neighboring Cdkn1a genes. EvoNC, a program for detecting selection in noncoding regions of nucleotide sequences, was used for this purpose (50). For protein coding sequences, the rate of nonsynonymous/synonymous substitution is used to detect the selection pressure and directionality of selection (i.e. positive or negative selection). Similar detection in noncoding sequences can be employed by calculating the rate of substitution relative to the rate of synonymous substitution in coding sequences using a parameter named ζ (please see Methods section for further details). A ζ value of 1 indicates that a site in a noncoding sequence evolved neutrally, whereas ζ > 1 and ζ <1 suggest positive and negative selection respectively (50). We concatenated the aligned domains A of lincRNA-p21 with the region of Cdkn1a exon for each ten mammals and analyzed the resulting sequences using EvoNC. A similar approach was undertaken to understand the evolution of lncRNA HOTAIR with respect to a neighboring protein coding gene (48). The program implemented three models, namely the neutral model, a two-category model, and a three-category model. For each model ζ was estimated as ζ0, ζ1 and ζ2. The results are shown in Table 4. The value of 4.94 found for ζ2 in the three-category case strongly suggested that the lincRNA-p21 region was under positive selection and evolved faster than Cdkn1a. Commonly, a gene with important biological function evolves slowly to maintain the role associated with it. However the exception can sometime be found when the gene is recent and still evolving (73), which is likely true for lincRNA-p21 gene. Nevertheless, the real factors that drive this positive selection are yet to be determined. Notably, in lncRNAs selection acts on structure rather than primary sequence that may also explain the rapid rate of evolution (23, 25, 74).

**Table 4.** Log-likelihood values and parameter estimates from EvoNC

| Method | K | $\omega$ | $\delta_0$ | $\delta_1$ | $\delta_2$ |
| --- | --- | --- | --- | --- | --- |
| Neutral | 3.02 | 0.29 | 0.14 | 1 |  |
| Two category | 2.59 | 0.29 | 0.5 | 1.75 |  |
| Three category | 3.02 | 0.29 | 0.14 | 1 | 4.94 |
| $\omega$ denotes rate of nonsynonymous/synonymous substitution in coding region. $\delta_0$, $\delta_1$ and $\delta_2$ are the rate of substitution in noncoding regions relative to the rate of synonymous substitution in coding regions for neutral, two-category, and three-category models. K refers to transition/transversion rate ratio |  |  |  |  |  |

### Structure prediction revealed two domains in lincRNA-p21 with invariable sequences and structures in mammals

LincRNA-p21 has been reported to interact with several proteins to exert its functions in cell. Therefore, it is important to identify the structure of functionally important domains in its sequence (28, 29). As long ncRNA sequences can adopt divergent structures in different species, determining the structure of the full-length lncRNA sequence experimentally is difficult and often unnecessary. Nevertheless, lncRNA may have a conserved structure in the form of functional domains. This feature prompted us to determine the sequence and structure of potentially functional and conserved regions instead of aiming structure prediction of the entire ~ 4kb lincRNA-p21 sequence.

Two constraints were used to facilitate the determination of the sequence and structure of possible functional domains. First, if a domain in the consensus structure of each query is essentially occupied by the sequences conserved in the ten mammals, then the domain may be considered as a functional domain. Second, functional domains should have invariant sequences and structures in all of the possible structures of the mammalian orthologs predicted by other method. Consensus covariance model for each query as shown in Figure 5A and Figure 6A, was configured through Infernal by aligning the identified orthologous sequences from different genomes that were previously obtained using Infernal’s structurebased genome searches (49). Table 5 shows the bit score and the average posterior probability (0 to 1) over all aligned nucleotides in each sequence of the alignment. High posterior probabilities in the range of 0.87 to 1 estimated for the models correspond to good confidence that the aligned nucleotide for each orthologous sequence belongs where it appears in the alignment. Structure of the individual orthologous sequence for both the queries was computed by PMmulti (51) and compared with the corresponding covariance model. Because each query produced only one high-scoring hit positioned between Srsf3 and Cdkn1a, we argue that the structures may be reasonable. Thermodynamic approach employed in Mfold (52) was used as other method to validate multiple potential structures for mammalian orthologous of lincRNA-p21.

**Figure 5.**
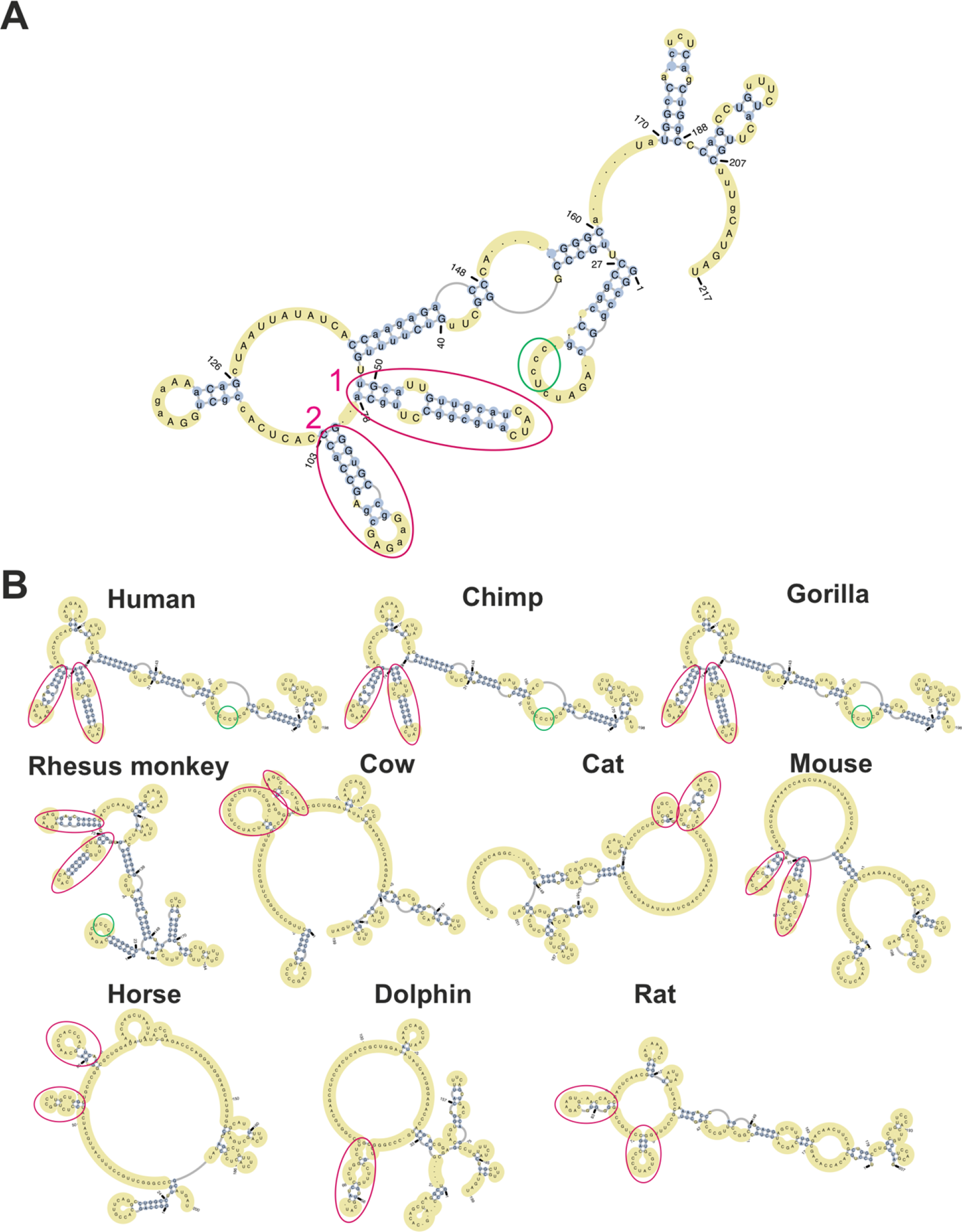
Predicted covariance model of domain A orthologous in mammals. (A) The consensus structure determined by Infernal. This consensus structure consists of two arcs and each arch with three substructures of hairpin loops. (B) The numbers of hairpin loops at the bottom vary from 1 (human) to 3 (dolphin) in different mammals as generated by PMmulti. Two hairpin loops marked as ‘1’ and ‘2’ in the top substructure of the consensus model that were found in all animals (except ‘2’ in dolphin) are marked in red ovals. Additionally, UCCC containing single stranded loop which is conserved in all primates are marked with a green circle.

**Figure 6.**
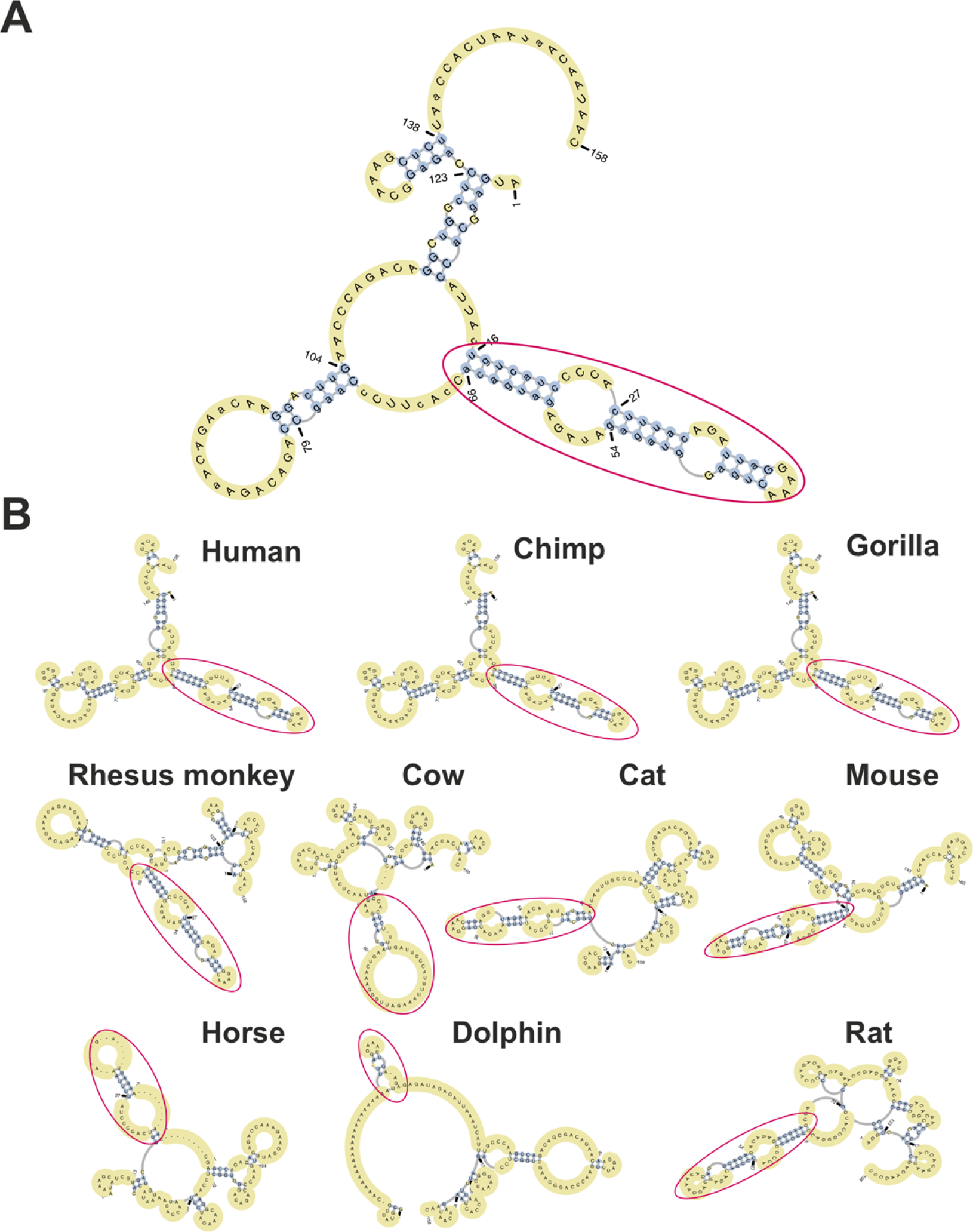
Predicted covariance model of domain B orthologous in mammals. (A) The consensus structure determined by Infernal. This consensus structure consists of an arc with three substructures of stem and loops. (B) The hairpin loop in the consensus model that was found in all animals is marked in red oval.

**Table 5.**
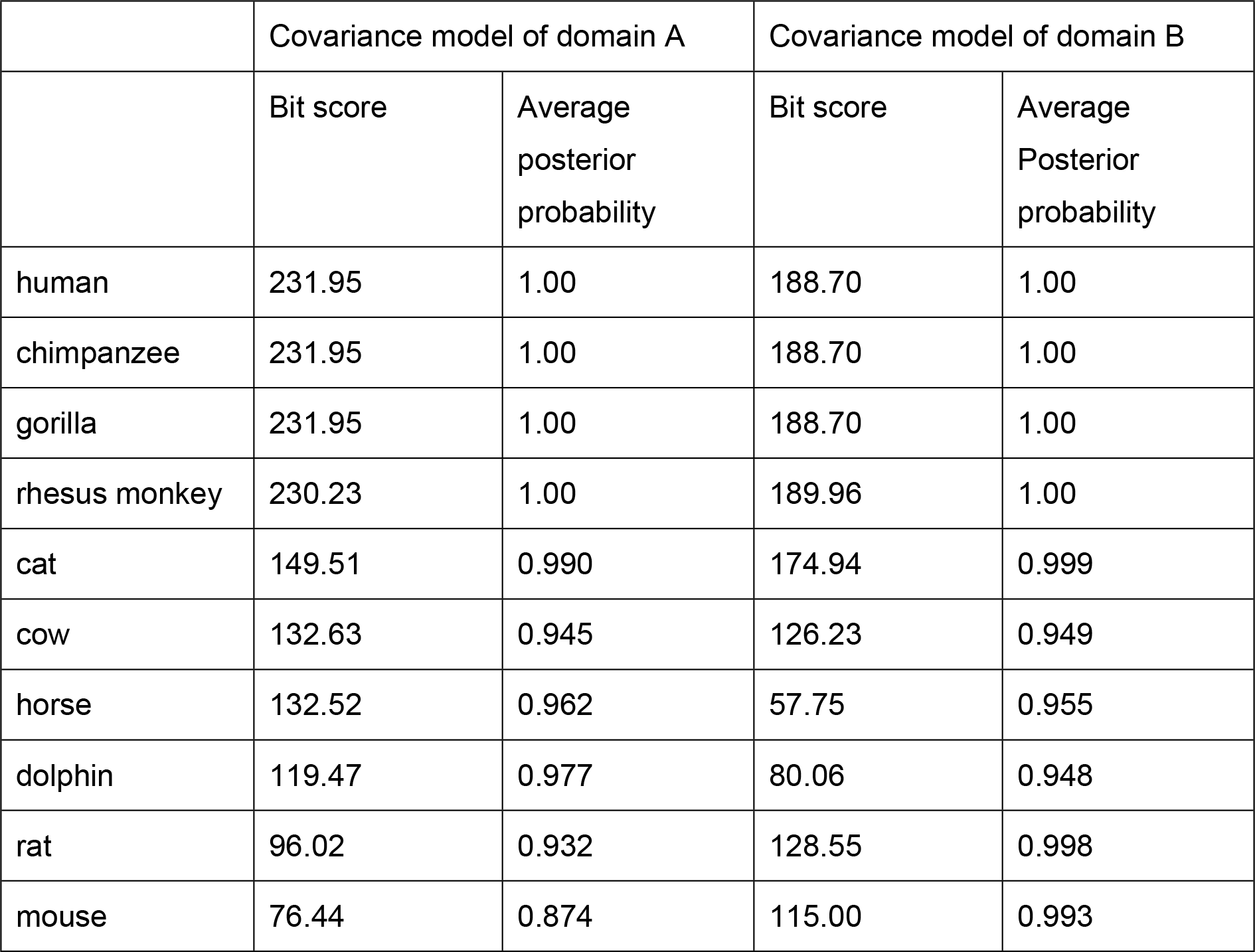
Alignment score of the orthologous sequences from different mammals to the consensus covariance model for domain A and domain B estimated from Infernal

|  | Covariance model of domain A |  | Covariance model of domain B |  |
| --- | --- | --- | --- | --- |
|  | Bit score | Average posterior probability | Bit score | Average Posterior probability |
| human | 231.95 | 1.00 | 188.70 | 1.00 |
| chimpanzee | 231.95 | 1.00 | 188.70 | 1.00 |
| gorilla | 231.95 | 1.00 | 188.70 | 1.00 |
| rhesus monkey | 230.23 | 1.00 | 189.96 | 1.00 |
| cat | 149.51 | 0.990 | 174.94 | 0.999 |
| cow | 132.63 | 0.945 | 126.23 | 0.949 |
| horse | 132.52 | 0.962 | 57.75 | 0.955 |
| dolphin | 119.47 | 0.977 | 80.06 | 0.948 |
| rat | 96.02 | 0.932 | 128.55 | 0.998 |
| mouse | 76.44 | 0.874 | 115.00 | 0.993 |

Figure 3 shows that the best structural hits were found as short fragments, one at the 5’ terminal region (domain A) and another towards the 3’ terminal region (domain B) of full-length lincRNA-p21. Because a 5’ region of lincRNA-p21 was previously shown to bind hnRNP-K (29), we assumed that the region is likely be conserved in mammals. Therefore, we attempted to identify a structured functional domain in domain A using the two constrains mentioned above. PseudoViewer program (56) was used to visualize the RNA secondary structure and it showed that the consensus covariance structure for domain A consists of two arcs, where each arch had three hairpin loops containing substructures (Figure 5A). These two arcs are connected by internal bulge containing central stem (Figure 5A). Two hairpin loops marked as ‘1’ and ‘2’ in the top substructure were found in all mammals (except in dolphin that lacks loop 2) (Figure 5B), which indicate that they could be functionally important sub-domains at the 5’ region of lincRNA-p21.

Further, we found a single occurrence of a UCCC sequence motif within the 5’ conserved domain (domain A) of lincRNA-p21. Notably, hnRNP-K has three RNA binding KH domains (KH1-3). KH domains in general have been shown to have specificity towards UCCC sequence containing motifs in DNA and RNA (75). In the consensus covariance structure, this sequence motif was found in region that has a high probability score to be a single stranded loop (Figure S1A). The UCCC motif remains conserved (for sequence and being single stranded) in the PMmulti predicted structures of lincRNA-p21 in all primates (Figure 5B). Therefore, we postulate that the UCCC containing region in domain A may constitute the binding motif for hnRNP-K. However, whether the flanking sequences have impact on protein binding can be investigated experimentally.

We next used Mfold to predict structures of orthologous sequences of domain A in each mammal (52). Mfold predicted 12 structures in human, 12 in chimpanzee, 12 in gorilla, 10 in rhesus monkey, 7 in cow, 11 in cat, 14 in dolphin, 19 in horse, 17 in mouse and 7 in rat. Representative structures from each mammal are shown in Figure S2. Notably, the first hairpin (marked as ‘1’ in Figure 5A) was found at the same position in 9 out of 12 predicted structures in human. A similar trend was observed for other mammals. Among them human, chimp, gorilla, rhesus monkey, horse, cat, mouse and rat contain a conserved ‘CAUC’ tetraloop in the single stranded hairpin loop (Figure S2) similar to hairpin loop of substructure ‘1’ in the Infernal predicted consensus structure of domain A (Figure 5A). In contrast, the substructure marked as hairpin loop ‘2’ in Figure 5A was found without any clear consensus substructure in the Mfold predicted structure (Figure S2).

Similarly, PseudoViewer showed that the consensus covariance structure for domain B consists of an arc with three substructures of stems and loops (Figure 6A and Figure S1B)). The stem-loop substructure marked in red oval was found in all animals (Figure 6B), which indicates that it could contain the functional domain at the 3’ terminal region of lincRNA-p21. As stated previously, this covariance structure was compared with all of the structures predicted by Mfold. Mfold predicted 2 structures in human, 2 in chimpanzee, 2 in gorilla, 5 in rhesus monkey, 5 in cow, 2 in cat, 1 in dolphin, 3 in horse, 8 in mouse and 4 in rat. Representative structures from each mammal are shown in Figure S3. Notably, the hairpin (red oval in Figure 6A) was found at the same position in the predicted structure in human (Figure S3). Similar results were obtained from all other animals except cow and horse (Figure S3). The loop in this hairpin structure consists of GAAA nucleotide sequence in human, chimp, gorilla, rhesus monkey, cat, dolphin, rat and mouse (Figure S3).

### Comparison of sequence and structures of the two postulated domains within full lincRNA-p21 predicted structure

Next, we predicted the secondary structures of the entire human lincRNA-p21 sequence using Mfold (52) and compared the sequence and structure of the two conserved fragments: domain A and domain B in full-length lincRNA-p21. The predicted sequence and structure of the domain A occurs near invariably in most structures. Mfold produced 19 structures for human full lincRNA-p21. The predicted sequence and structure of the fragment ‘1’ in domain A occurs in 18 of 19 full-length lincRNA-p21 structures and the predicted structure of fragment ‘2’ in domain A occurs in 8 out of those 18 structures that include the lowest free energy full-length human lincRNA-p21 structure (encircled in red, Figure S4). Similarly, predicted structure of the fragment with the ‘GAAA’ tetraloop motif in domain B occurs in 8 of 19 full lincRNA-p21 human structures (encircled in red, Figure S4).

### Alu repeats exits in human but show poor conservation in mammals

Isoforms of human lincRNA-p21 were found to contain inverted repeat Alu elements (IRAlus) (76). The sense element of these IRAlus is located at positions 2589-2895 and the antisense Alu element is located at positions 1351-1651. These IRAlus were shown to form independent structural domains in the context of full human lincRNA-p21 (76). IRAlus formed by human lincRNA-p21 were found to be important regulator for its cellular localization over the course of the stress response. We searched several vertebrate genomes in the UCSC databases for matches to human lincRNA-p21 IRAlus using BLAT (62). The result depicted in Figure S5 shows that the close matches for both the IRAlus were found only in primates (Figure S5A and S5B). BLAT searches produced hits in chimpanzee, gorilla (partial) and rhesus monkey in between Srsf3 and Cdkn1a genes. However, no significant hits were found in non-primate mammals and other vertebrates. This is in contrast to conserved domains A and B in the 5’ and 3’ terminus respectively of lincRNA-p21 that were found to exist in all mammalian orthologs (Figure S6A and S6B). Therefore, the IRAlu regions seem to have undergone recent evolution in lincRNA-p21 imparting further functions to the RNA.

## Conclusions

Since, orthologous sequence of lincRNA-p21 identified using the RNA homology search software Infernal contain sequence mismatch with gaps, we inferred that lincRNA-p21 harbor poorly conserved sequences but considerably conserved structures in ten examined mammals, a feature that is prevalent in other lncRNAs (65). Infernal search found just one high scoring hit in each mammal located between Cdkn1a and Srsf3 genes. Thus, it can be concluded that full length lincRNA-p21 exists only in mammals. Additionally, except for one high scoring hit, several low-scoring hits were produced in many other places in mammalian and other vertebrate genomes, which suggests there may exist other lncRNAs that share similar functional domains with lincRNA-p21. However, the extent of conservation in lncRNAs to preserve the function with simultaneously evolving sequences remains elusive.

Phylogenetic analysis of conserved segments of lincRNA-p21 in ten mammals covering primates, rodents, carnivores and ungulates revealed discrete evolutionary dynamics for orthologous sequences of lincRNA-p21 with different nucleotide substitution rates between clades. Notably, both the domains were found to evolve in a slow rate in primates than in non-primate mammals suggesting for strong functional constrains which may restrict further evolution of lincRNA-p21 conserved domains in primates. Comparison between lincRNA-p21 orthologous sequences of domain A and the exon of Cdkn1a (a protein coding gene) clearly showed that the lincRNA-p21 underwent positive selection and evolved significantly faster than the neighboring Cdkn1a exon. The facts that lincRNA-p21 is not found in non-mammalian vertebrates and has evolved faster than it’s nearby genes suggested that it is a recent gene. Given that most lncRNAs, including Xist and HOTAIR, have so far only been found in mammals, it is interesting question to ask as when and why these lncRNAs emerged in higher vertebrates to mediate genome modifications and other functions.

A comparative computational approach was undertaken to predict the sequence and structure of conserved functional domains of lincRNA-p21 in mammals. PMmulti (51) and Mfold (52) were used to predict multiple potential structures for orthologous of lincRNA-p21 domains in ten mammals. Two consecutive stem-loops in domain A at 5’ terminal region and a single stem-loop in domain B at 3’ terminal region of lincRNA-p21 were postulated through structure prediction where their sequence and structure invariably appeared in several predicted structures of ten different mammals, also in their consensus model along with full lincRNA-p21. This also suggest that these structural fragments located at the 5’ end and close to the 3’ end of lincRNA-p21 could be functional domains of the lincRNA-p21. The recently identified IRAlus in lincRNA-p21 seems to have evolved recently in human, suggesting a still continuing evolution of lincRNA-p21.

Producing experimental data to determine structures of lncRNAs is challenging and time-consuming. Hence, the results of our structure prediction should be useful for further experimental studies on determining the structure of lincRNA-p21 and its interaction with protein partners. A similar strategy can be extended to study structure and functions of other lncRNAs.

## Abbreviations

lincRNA-p21, lncRNA, MFOLD, BLAST, Infernal, IRAlus

## Author Contributions

M.S. conceived the research; M.S. and A.M. designed the research; A.M. performed the research; M.S. and A.M. analyzed the results and wrote the manuscript.

## Funding

This work is supported by a Early Career Fellowship from Wellcome Trust/ DBT India Alliance to A.M. (IA/E/15/1/502321). Research in M.S. lab is supported by grants from Department of Biotechnology (DBT), India (BT/PR15829/BRB/10/1469/2015) and IISc-DBT partnership program.

## Acknowledgements

The authors also acknowledge funding for infrastructural support from the following programs of the Government of India: DST-FIST, UGC-CAS, and the DBT-IISc partnership program. M.S. is a recipient of Ramalingaswami Fellowship.

## Competing Interests

The authors declare that there are no competing interests associated with the manuscript.

